# Single-molecule magnetic tweezers reveals that TAV2b-derived peptides underwind and stabilize double-stranded RNA

**DOI:** 10.1101/2024.09.05.611413

**Authors:** Zainab M. Rashid, Salina Quack, Misha Klein, Quinte Smitskamp, Pim P. B. America, Marvin A. Albers, Jannik Paulus, Tom N. Grossmann, David Dulin

## Abstract

Double-stranded RNA (dsRNA) has evolved into a key tool in understanding and regulating biological processes, with promising implications in therapeutics. However, its efficacy is often limited due to instability in biological settings. Recently, the development of peptidic dsRNA binders derived from naturally occurring RNA-binding proteins has emerged as a favorable starting point to address this limitation. Nevertheless, it remains unclear how these high affinity dsRNA binders alter the structure and flexibility of dsRNA. To this end, we employed single-molecule magnetic tweezers experiments to investigate the effects of TAV2b-derived peptidic dsRNA binders on the mechanical properties of dsRNA. Torque spectroscopy assays demonstrated that these peptides underwind dsRNA, while also stabilizing the duplex. Additionally, force spectroscopy experiments demonstrate that a wild type TAV2b peptide derivative extends the contour length and lowers the bending rigidity of dsRNA, while a homodimeric version triggers the formation of higher order complexes at forces below 1 pN. Our study presents a quantitative approach to investigate how these peptides alter the structure of dsRNA, and whether peptide structural design alters the affinity to dsRNA and its stability. This approach could inform the design of more potent and effective dsRNA binders in the efforts to advance RNA therapeutics.

## Introduction

RNA has emerged as a pivotal tool in the understanding and modulation of biological processes (1, 2). One example involves small interfering RNAs (siRNAs) which function as potent gene silencers by inducing post-transcriptional RNA interference, thereby downregulating or preventing the expression of target proteins (3–5). siRNAs comprise short double-stranded RNA (dsRNA) that demonstrate promising therapeutic potential. However, their therapeutic efficacy is limited by low stability of RNA in the biological context (2, 5, 6). To enhance the stability, the dsRNA can be chemically modified or enclosed in nanocarriers (3, 5). The functionality of this approach, however, can be limiting as the modifications may either disrupt RNA activity or are optimized for a specific application (7, 8).

A promising strategy to modulate dsRNA stability and function involves the use of peptides derived from naturally occurring RNA-binding proteins (9–11). For example, dsRNA-specific peptide binders have been derived from the tomato aspermy virus 2b (TAV2b) protein of the *Bromoviridae* family of plant viruses (8, 12–14). TAV2b functions as a gene silencing suppressor by binding to dsRNA, thereby counteracting host defense mechanisms (15). This protein primarily consists of two alpha helices and binds to the major groove of dsRNA as a dimer, supported by an N-terminal leucine zipper-like dimerization motif (**Figure 1A**). It binds to dsRNA with high affinity and in a sequence-independent manner (15). Initial studies used a 33-mer TAV2b-derived peptide (wt33, **Figure 1B**) which represents the core RNA interaction motif and retained moderate dsRNA binding affinity (13). To introduce an environment-sensitive dsRNA release mechanism, a homodimeric peptide 1’-1’ was designed. It only contains helix 1, dimerized via a disulfide bond, replacing the leucine zipper-like motif present in the native protein (**Figure 1AC**). Upon exposure to the reducing conditions found in the cytosol the disulphide bond gets cleaved, thereby releasing dsRNA. Importantly, peptide 1’-1’ also retained its affinity for dsRNA (8). Taken together, both peptides, wt33 and 1’-1’, bind specifically to dsRNA and stabilize their duplex structure (8, 13).

**Figure 1:**
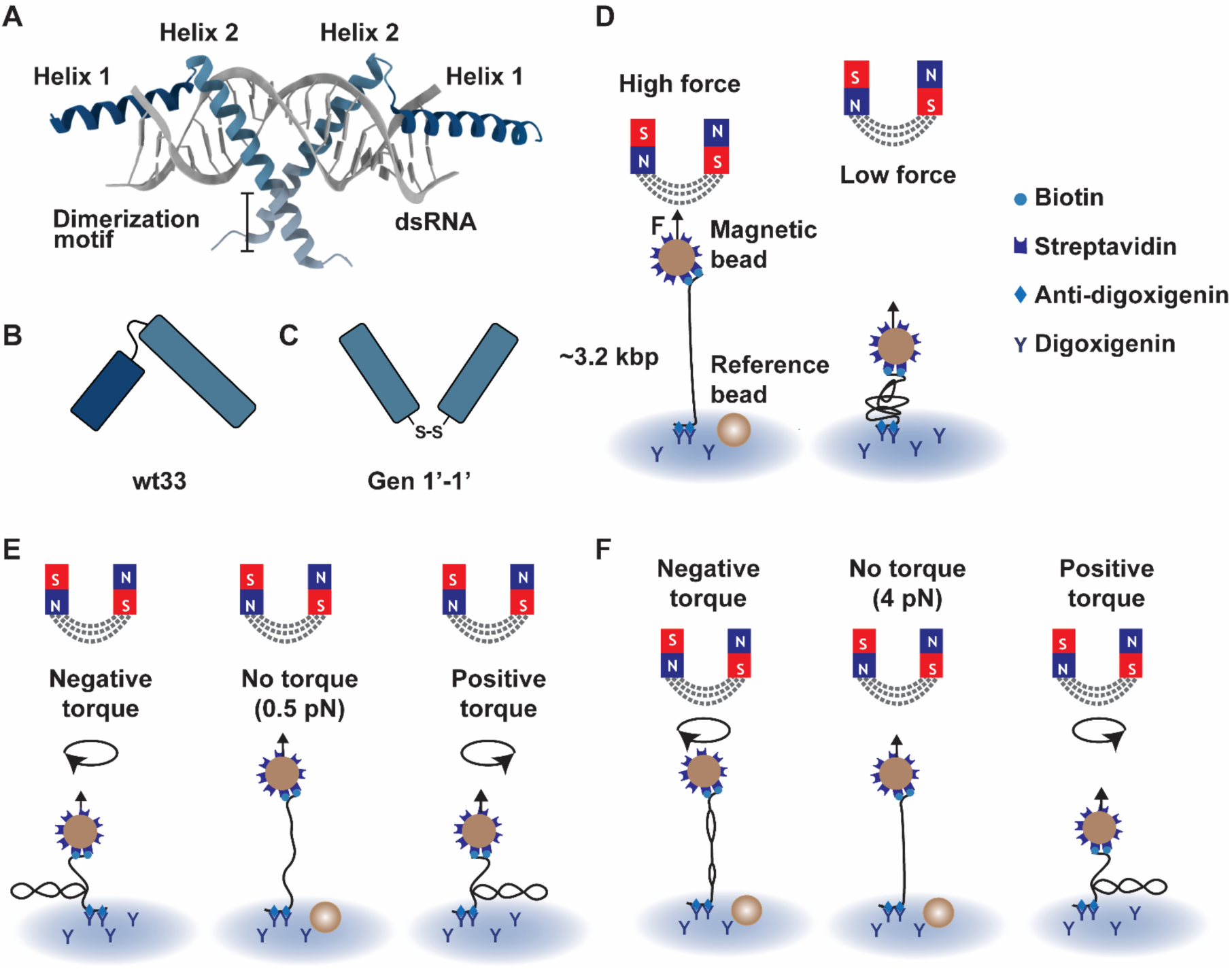
Force- and torque-spectroscopy of TAV2b derivatives in interaction with dsRNA using magnetic tweezers. **(A, B, C)** Schematics of TAV2b interactions with dsRNA and its derivatives (B) wt33 (aa 21-53) (13) and (C) 1’-1’ (aa 18-37) (8). Figure inspired by McLoughlin et al. 2023. **(D, E, F)** Schematic of the force-extension and rotation-extension experiments, respectively, in a magnetic tweezers assay.

While these studies have demonstrated that RNA-binding proteins can serve as starting points for the design of peptidic dsRNA binders to stabilize dsRNA, insights into their impact on the mechanical properties of dsRNA and therefore the modulation of RNA structure and flexibility are lacking. Here, we employed real-time, high-throughput magnetic tweezers (MT) to investigate how peptides wt33 and 1’-1’ impact the mechanical properties of dsRNA. Using torque spectroscopy experiments, we show that wt33 and 1’-1’ underwind and stabilize the dsRNA upon binding. Additionally, the force spectroscopy experiments showed that upon binding, peptide wt33 increases the dsRNA tether contour length by ∼10% and significantly lowers its persistence length compared to bare dsRNA. Peptide 1’-1’ extends the dsRNA but compacts it to form higher order complexes at forces lower than 1 pN. Our study presents a quantitative approach to analyze how novel peptidic dsRNA binders modulate the flexibility, length, stability, and natural twist of dsRNA. These findings can complement bulk biochemical assays, providing further insights into how peptide design may impact dsRNA structure and consequently the efficacy of such dsRNA binders. This approach informs the development of more potent and effective dsRNA binders in the efforts to advance RNA therapeutics.

## Results

### Investigating mechanical properties of dsRNA-peptide interactions at the single-molecule level

We designed a high-throughput magnetic tweezers assay to monitor changes in contour length, persistence length and twist of dsRNA following the addition of TAV2b peptide derivatives (16). A ∼3.2 kbp dsRNA is flanked with a biotin-labelled handle on one end and a digoxigenin-labelled handle on the other end. It is tethered between a streptavidin-coated magnetic bead and an anti-digoxigenin coated flow cell surface (**Figure 1D**) (16–19). Multiple attachment points at both ends render the dsRNA tether coilable (**Figure 1E**). The pair of permanent magnets located above the flow cell apply an attractive force to the magnetic bead, stretching the dsRNA tethers (16). The attractive force exerted is controlled by the magnet’s distance to the bead (**Figure 1D**) (16). To track the extension of the dsRNA tether as a function of force, the height of the magnet is continuously varied in three consecutive measurements, referred to here as “dynamic force extensions” (20). Furthermore, rotation of the magnets, and consequently the magnetic bead, enables torque spectroscopy of the dsRNA at constant force (**Figure 1EF**, **Material and Methods**) (16). To probe the change in twist of the dsRNA-peptide complexes, rotation experiments can be performed on coilable dsRNA at 4 pN and 0.5 pN, here referred to as “rotation-extension” (**Figure 1EF, Material and Methods**). A C++/CUDA bead tracking software integrated in LabView tracks the tri-dimensional positions (x, y, and z) of hundreds of beads in real-time at a nanometer resolution (**Materials and Methods**) (16, 21).

### Peptide wt33 underwinds and alters the contour and persistence length of dsRNA

We first employed dynamic force-extensions to extract changes in the contour and the persistence length of dsRNA upon addition of 10 μM of wt33 at (**Figure 2A, S1A**). The non-extensible worm-like chain (WLC) model was fitted to each individual trace for both dsRNA and the dsRNA-peptide complex (**Figure 2A, Materials and Methods**) (20, 22). The WLC fits yielded an average contour length of *L*_*C*_ = (968 ± 75) *nm* for dsRNA, comparable to the expected crystallographic length of a ∼3.2 kbp dsRNA (**Figure 2B**, **Table S1**) (23, 24). In the presence of wt33, the contour length increased to *L*_*C*_ = (1092 ± 94) *nm*, suggesting that the peptide extends the dsRNA upon binding (**Figure 2B, Table S1**). Furthermore, the persistence length showed a pronounced change upon addition of 10 μM wt33, with a measured value of *L*_*P*_ = (10 ± 2) *nm* (**Figure 2C**). This is significantly lower than the persistence length of dsRNA, reported as *L*_*P*_ = (46 ± 9) *nm*, which is in approximate agreement with previous measurements (23, 24). Peptide wt33 therefore significantly reduces the bending rigidity of dsRNA upon binding (**Table S1**).

**Figure 2:**
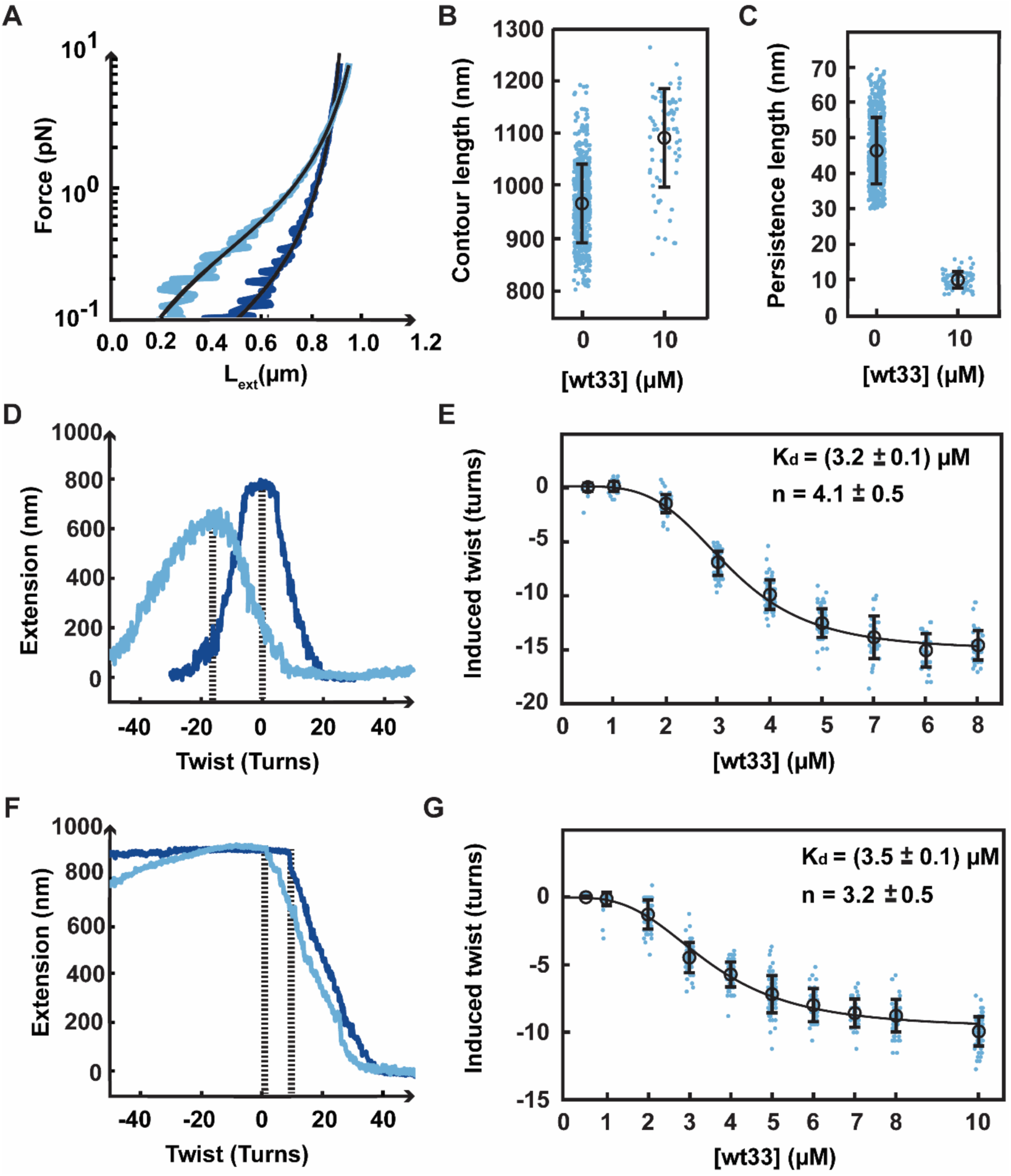
Peptide wt33 decreases the persistence length and unwinds dsRNA upon binding. **(A)** Force-extension experiments for a torsionally unconstrained dsRNA tether before (dark blue) and after (light blue) the addition of 10 µM wt33 peptide. The black solid line is a WLC model fit to the data. **(B)** Plot displaying the distribution of contour length for torsionally unconstrained dsRNA at 10 µM of wt33. **(C)** Plot showing the decrease in persistence length for torsionally unconstrained dsRNA at 10 µM of wt33. **(D)** Rotation-extension curves for a torsionally constrained dsRNA alone (dark blue) and in presence of 8 µM of wt33 (light blue) at 0.5 pN. **(E)** Zero-twist peak position as a function of wt33 concentration measured at 0.5 pN. **(F)** Rotation-extension curves for a torsionally constrained dsRNA alone (dark blue) and in presence of 8 µM of wt33 (light blue) at 4 pN. **(G)** Zero-twist peak position as a function of wt33 concentration measured at 4pN. (**D,F**) The zero twist positions for both traces are indicated with a dashed line. (**E, G**) The black solid line is a Hill model fit to the mean positions (circles) (**Materials and Methods**). The blue dots represent the values extracted for each measurement, the black circles are the mean values, and the error bars represent one standard deviation.

To test whether wt33 binding to dsRNA changed its twist, we performed rotation-extension experiments on coilable dsRNA at different concentrations of wt33 (**Figure S1B**, **Table S2**) (16). At low force, the extension of bare dsRNA decreases symmetrically for both positive and negative turns (**Figure 1E**). At 4 pN, the extension is constant for negative turns, as dsRNA unwinds, and decreases for positive turns once the buckling transition to form plectonemes is passed (**Figure 1F**) (16, 24–26). We could reproduce these results for dsRNA at 0.5 and 4 pN (**Figure 2D-G**) (13, 18, 19). Upon addition of wt33, the zero-turn peak position shifted to the left, i.e. negative turns, which suggests that the peptide underwound dsRNA upon binding (**Figure 2D**). We also noted that the maximum extension observed for the dsRNA-peptide complex is lower compared to that of bare dsRNA (**Figure 2D**), which is due to a decrease in persistence length of the tether (**Figure 2C**). Furthermore, we observe that the rotation curve broadens and the slope of the plectonemic regions decrease upon addition of wt33 (**Figure 2D**). This behaviour can be qualitatively explained using a simplified elastic model on DNA mechanics. The model predicts that the size of the supercoil in the plectonemic region scales as the square root of the persistence length: Δ*z ∼* √*L_p_*. Given that the persistence length significantly decreases upon addition of wt33 (**Figure 2C**), this model predicts a reduced Δ*z* in the plectonemic region, thereby resulting in a decreased slope (25, 26).

Moreover, we observe asymmetrical slopes in the plectonemic regions during application of positive and negative torque in the presence of wt33, i.e. (0.04 ± 0.01) µ*m*/*turn* vs (0.02 ± 0.01) µ*m*/*turn*, respectively. This suggests a torque dependent binding mechanism (**Figure 2D**). The slope of the plectonemic region during application of negative torque is gentler compared to that observed under positive torque. We can use the simple elastic model previously described to interpret this behaviour. Applying negative torque enhances binding of wt33 to dsRNA, which subsequently reduces the persistence length. This decrease in persistence length results in a smaller Δ*z* in the plectonemic region, thereby leading to a decrease in slope. In contrast, positive torque would disrupt binding, and result in a comparatively steeper slope. Our data shows that wt33 underwinds dsRNA, hence it is plausible that underwinding of dsRNA promotes wt33 binding to dsRNA.

We observed that the zero-twist peak shifted from 0 to -15 turns at 0.5 pN, saturating above 6 µM (**Figure 2E**). The mean position of the zero-twist peak is well described by a Hill model, with a *K*_*d*_ = (3.2 ± 0.1) µ*M* and *n* = 4.1 ± 0.5, i.e. much larger than 1 and therefore indicating a highly cooperative association of wt33 to dsRNA (**Figure 2E**, **Table 1**). At 4 pN, we monitored the shift of the buckling transition upon positive supercoiling, i.e. the number of turns at which the coilable tether starts forming plectonemes (**Figure 2F**). Fitting a Hill model to the buckling transition, we extracted a *K*_*d*_ = (3.5 ± 0.1) µ*M* and *n* = 3.2 ± 0.5 (**Figure 2G**, **Table 1**), which is in good agreement with the values obtained at 0.5 pN (**Figure 2E**). Additionally, we notice a decrease in the end-to-end distance during application of negative supercoil. This indicates that the critical torque required to unwind dsRNA has increased, suggesting that the peptide stabilizes the dsRNA structure upon binding (**Figure 2F**).

**Table 1:** Fitting parameters using the Hill equation for peptide wt33 and 1’-1’.

| Peptide | $(K_d \pm 2SE) \mu M$ | Delta twist $\pm 2SE$ | Hill coefficient $\pm 2SE$ |
| --- | --- | --- | --- |
| wt33 (0.5 pN) | $3.2 \pm 0.1$ | $-15.0 \pm 0.2$ | $4.1 \pm 0.5$ |
| wt33 (4 pN) | $3.5 \pm 0.1$ | $-9.7 \pm 0.2$ | $3.2 \pm 0.5$ |
| 1'-1' (4 pN) | $1.5 \pm 0.1$ | $-13.1 \pm 0.2$ | $3.1 \pm 0.2$ |

### Dimeric peptide 1’-1’ underwinds dsRNA and dissociates in reducing conditions

Using the same experimental approach, we next investigated how 1’-1’ altered the mechanical properties of dsRNA compared to wt33. Additionally, we evaluated its binding affinity to dsRNA relative to wt33. To probe changes in the contour and persistence length of dsRNA, we performed a dynamic force-extension experiment as described for peptide wt33. We observed that upon decreasing the applied force to below 1 pN, the tethers collapsed. In this regime, thermal fluctuations enable long distance interactions between peptides associated to the dsRNA (**Figure 3A**, **Figure S2A**). Indeed, we have shown that 1’-1’ forms higher order structures in the presence of dsRNA (8). Consequently, we were unable to fit a WLC model on such traces to extract the changes in the contour and persistence length.

**Figure 3:**
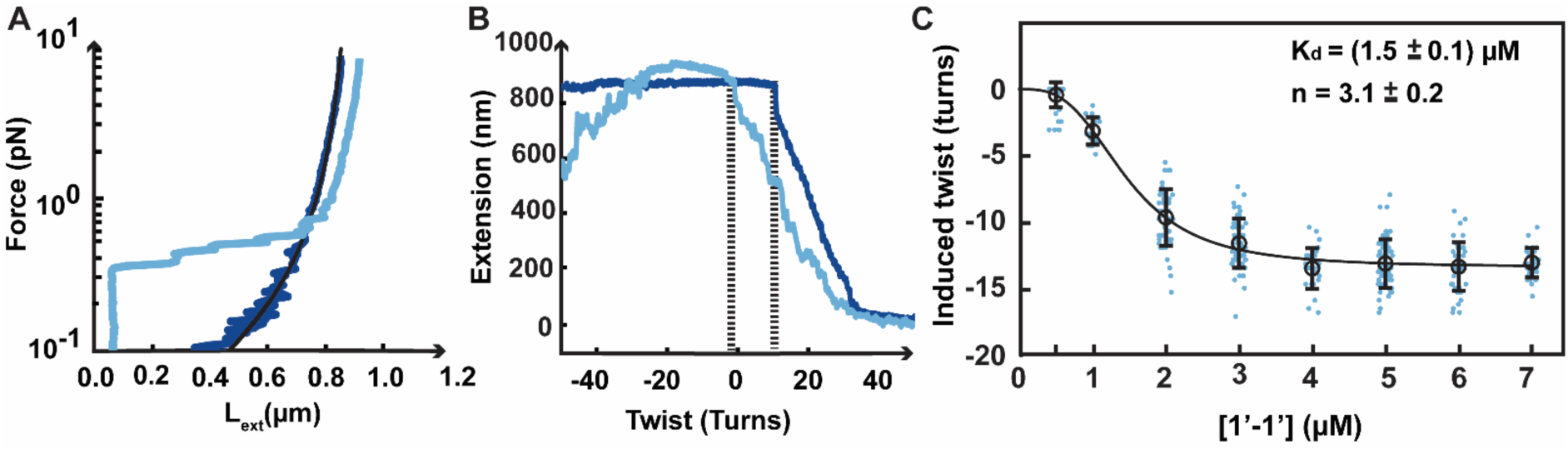
Peptide 1’-1’ unwinds dsRNA upon binding. **(A)** Force-extension experiments for a torsionally unconstrained dsRNA tether before (dark blue) and after (light blue) the addition of 7 µM 1’-1’ peptide. The black solid line is a WLC model fit to the data. **(B)** Rotation-extension experiment for a torsionally constrained dsRNA in absence (dark blue) and in the presence of 7 µM 1’-1’ peptide (light blue). The measurements were performed at 4 pN. The respective zero-twist peak positions are indicated by a dashed line. **(C)** Zero-twist peak position as a function of 1’-1’ concentration. The black solid line is a Hill model fit to the mean positions (circles) (**Materials and Methods**). The blue dots represent the values extracted for each measurement, the black circles are the mean values and the error bars represent one standard deviation.

To determine the change in dsRNA twist induced by 1’-1’ binding, we performed rotation-extension experiments at varying concentration of 1’-1’ peptide. As the tether collapsed at low force (**Figure S2B**) (8), we performed these experiments at 4 pN (**Figure 3B**). Upon addition of 1’-1’, the rotation-extension curve is left-shifted, suggesting that this peptide also underwinds dsRNA upon binding (**Figure 3B**). Furthermore, we notice a decrease in the end-to-end distance during application of negative turns, suggesting that the peptide stabilizes the dsRNA structure upon binding. To evaluate and compare the dissociation constant of peptide 1’-1’ to wt33, we performed a concentration sweep and monitored how the buckling transition point shifts for 1’-1’ (**Figure 3C**). With increasing peptide concentrations, the dsRNA tethers were progressively unwound and plateaued at about 4 μM, with a -13 turn shift with respect to bare dsRNA (**Table S3**). Fitting the Hill model on the mean buckling transition shift values yielded a *K*_*d*_ = (1.5 ± 0.04) µ*M*, further supporting the improved affinity in respect to peptide wt33 (8). We also extracted a Hill coefficient of *n* = 3.1 ± 0.2 suggesting that 1’-1’ peptide associates to dsRNA in a highly cooperative fashion (**Table 1**).

Furthermore, to mimic the reducing conditions present in the cytosol, that trigger release of the RNA from the peptide, we performed a dynamic force-extension assay in the presence of tris(2-carboxyethyl) phosphine (TCEP). Our experiments show that cleavage of the disulfide bond upon exposure to reducing conditions, renders peptide 1’-1’ ineffective in binding to dsRNA. This is shown by the absence of higher-order structure formation at forces below 1 pN, and a force-extension behaviour identical to that of dsRNA following treatment with (TCEP) (**Figure S3**, **Materials and Methods**). This indicates that the disulfide bond functions as an effective dsRNA release mechanism upon exposure to reducing conditions, such as in the cytosol (8).

## Discussion

The increased use of dsRNA in biotechnological and biomedical applications is paralleled with efforts directed at stabilizing dsRNA for increased efficacy and understanding the effects of peptide and protein binding. Although bulk biochemical studies have shown that RNA-binding proteins can serve as suitable starting points for the design of peptidic dsRNA binders, we lack quantitative data on how these peptides modulate the structure and flexibility of dsRNA. To address this gap, we used single-molecule magnetic tweezers to investigate how TAV2b-derived peptides, wt33 and 1’-1’, alter the twist, persistence and contour lengths of dsRNA.

Overlaying the TAV2b-dsRNA crystal structure (pdb:2zi0) with an unbound dsRNA of an identical sequence showed that upon binding to the major groove, the TAV2b dimer underwinds the dsRNA helix (**Figure 4A-D**). This is indicated by an increase in the number of base pairs per helical turn, and a widened major groove in the presence of TAV2b. A-form dsRNA contains 11 bp per helical turn and a major groove distance of 11 Å, whereas TAV2b-bound dsRNA has 12 bp per helical turn and a major groove distance of 19 Å **(Figure 4A-D**, **Materials and Methods)**. This structural alteration can also be observed with our torque-based experiments, which demonstrated that the derived peptides retain the ability to unwind dsRNA (**Figure 4E**). Using the average maximum turns recorded at saturating peptide concentrations and the peptide binding footprints, the degree of unwinding induced by each peptide can be quantified. For peptide wt33, which has a binding footprint of 11 base pairs, the torque assays at 0.5 pN showed an average of ∼-15 turns at saturated peptide conditions. Assuming full occupancy of the dsRNA tether at saturated peptide conditions, a ∼3.2 kbp tether would be bound by 290 peptides. If each peptide contributes equally to the unwinding, a single peptide unwinds dsRNA by -0.05 turns, corresponding to an induced twist of approximately -18.5°. However, torque assays performed at 4 pN under saturating peptide concentrations resulted in an average twist of about -10 turns. Under these conditions, each peptide induces a twist of -0.03 turns, corresponding to -12.4° per peptide. Furthermore, for peptide 1’-1’, the torque experiments at 4 pN showed an unwinding of ∼-13 turns at saturating peptide concentrations. Using the same assumptions and given that the binding footprint is identical to wt33, our data suggests that each peptide unwinds dsRNA by -0.04 turns, corresponding to a twist of -16°. Moreover, application of negative torque at 4 pN demonstrated that these peptides can stabilize the structure of dsRNA, consistent with bulk experiments performed with these peptides. Overall, we showed that this approach provides quantitative structural insights into how these peptides alter and stabilize the helical structure of dsRNA, which can be complemented with crystallographic studies and bulk biochemical assays.

**Figure 4:**
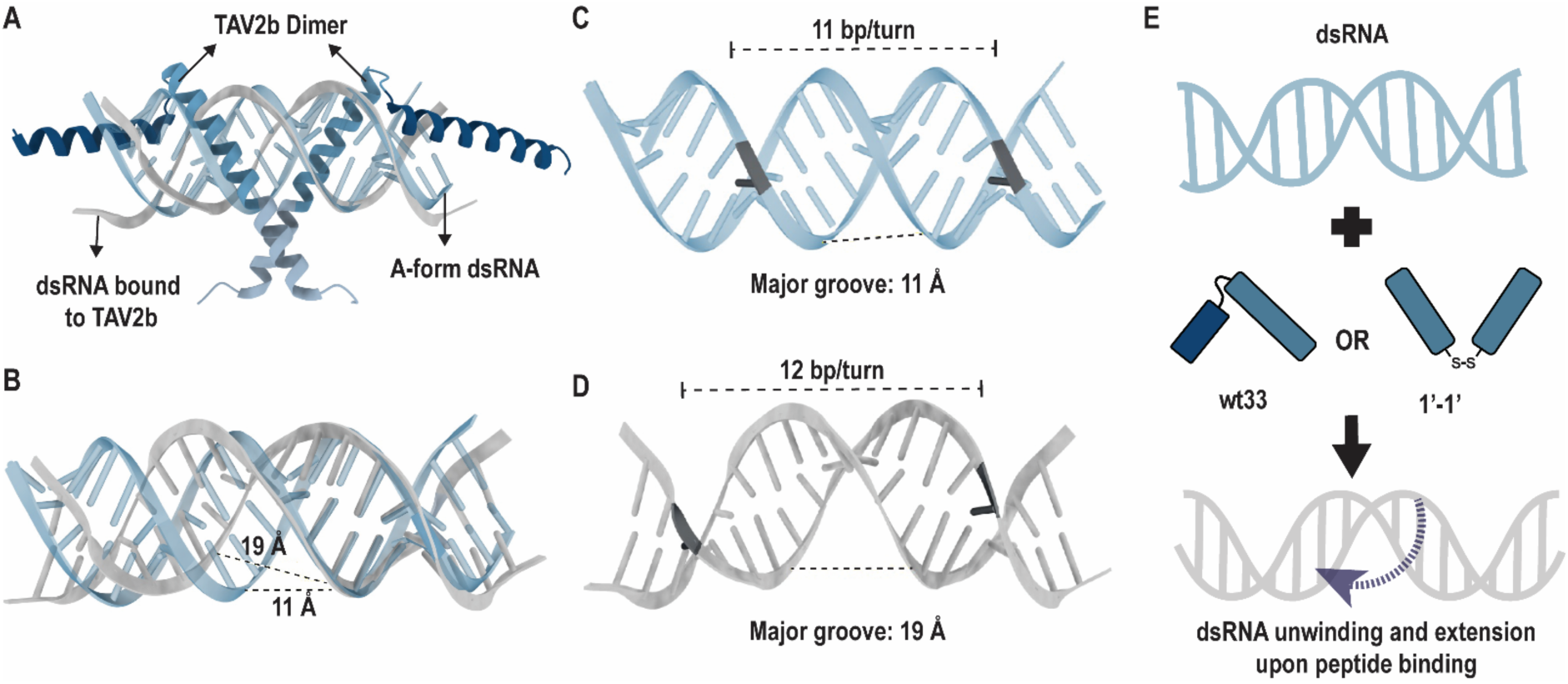
Structural analysis of dsRNA underwinding by TAV2b. **(A)** TAV2b dimer-dsRNA crystal structure (pdb: 2zi0) superimposed with free A-form dsRNA (blue) (**Materials and Methods)**. **(B)** A-form dsRNA (blue) aligned with TAV2b-bound dsRNA (grey) illustrating a widened major groove in presence of TAV2b. **(C)** Structure of A-form dsRNA highlighting number of base pairs per helical turn and distance of the major groove. **(D)** Structure of TAV2b-bound dsRNA highlighting increased number of base pairs per helical turn and widened major groove. **(E)** Model illustrating dsRNA underwinding in the presence of peptide wt33 and 1’-1’.

Implementing the Hill model to describe the shift in the zero-twist peak position with increasing peptide concentrations demonstrated that peptide 1’-1’ has a higher affinity to dsRNA relative to peptide wt33, suggested by a lower dissociation constant. This data further substantiates that helix 1 is sufficient for effective dsRNA binding when present as a dimer (8) and highlights the importance of structural assembly for dsRNA recognition. Moreover, bulk biochemical assays have established that stabilizing the RNA-binding helices by employing hydrocarbon staples results in improved RNA duplex stability in a dsRNA-peptide complex (8). To further investigate this finding, our torque assay can also assess whether this structural modification enhances its affinity to dsRNA and alters the dsRNA helical structure differently compared to other peptides. We can further investigate if it improves stabilization of the A-form dsRNA, which would be observed as a more pronounced change in the extension upon application of the negative torque at 4 pN. These studies could inform and inspire the structural design of more efficient RNA-binding peptides, potentially improving dsRNA delivery for biotechnological or biomedical applications.

Additionally, the Hill model yielded a Hill coefficient (n) greater than 1 for both peptides, indicating cooperativity in binding to dsRNA. This coefficient, derived from the mass action law, reflects the number of peptides required to establish an interaction with the dsRNA tether. However, it only represents the binding cooperativity to dsRNA and does not quantify the interaction strength between the peptides. For the torque experiments, using the Hill equation treats the ∼3.2 kbp dsRNA molecule as a single substrate, despite being able to accommodate up to ∼290 peptides. Therefore, instead of interpreting the Hill coefficient as the number of peptides required to interact with the dsRNA tether, we consider it as an indication that a bound peptide quite strongly attracts another peptide to bind next to it. Overall, the Hill coefficient values suggest that this process is cooperative.

The force spectroscopy experiments have demonstrated that peptide wt33 significantly reduces the bending rigidity and increases the contour length of dsRNA, due to helix underwinding. A quantitative comparison to peptide 1’-1’ was not possible because the tether collapses at lower forces and forms higher order structures. However, the dynamic-force extension and rotation-extension experiments indicate an increase in dsRNA extension in the presence of the peptide 1’-1’, suggesting that, like wt33, it retains the ability to extend dsRNA due to helix underwinding (**Figure 3A-B**). Additionally, given that these peptides are often engineered for therapeutic applications, such as delivery of siRNA to the cells, it is crucial to further study how these peptides are processed in the cell following RNA delivery and whether these peptides would trigger an immune response given their foreign origin. This understanding is essential for designing effective and safe therapeutic tools.

Altogether, this study uses a single molecule approach to quantitatively assess how high-affinity dsRNA-binders impact the mechanical properties and helical structure of dsRNA. It offers insights into how the structural design of these peptides influences dsRNA stability, and the binding affinity to dsRNA. In addition to complementing bulk biochemical assays, this method provides an innovative approach to inform the development of potent and effective dsRNA binders, equipped with spatiotemporal dsRNA release mechanisms. Ultimately, this potentially accelerates developments in targeted RNA delivery, personalized medicine and drug formulation.

## Materials and Methods

### Double-stranded RNA (dsRNA) synthesis

A detailed approach on dsRNA synthesis has been provided in Ref. (17). In short, a pBAD plasmid was used to generate DNA fragments using Phusion polymerase and primers containing a T7 promoter. Next, ssRNA fragments were obtained using in vitro transcription (HiScribe Kit, New England Biolabs) and were mono phosphorylated using 5’-Polyphosphatase (Biosearch Technologies). Lastly, the ssRNA fragments were annealed to form dsRNA and ligated to handles containing Digoxigenin-UTP (Jena Biosciences) and Biotin-UTP (Jena Biosciences) using T4 RNA ligase 2 (New England Biolabs).

### Peptide synthesis

An extensive protocol on synthesis of peptide wt33 and 1’-1’ has been described in Ref. (8, 13). The peptides were dissolved in DMSO to a concentration of 10 mM and stored at -20 °C. The aliquots for peptide wt33 and 1’-1’ were thawed shortly before the experiments. The peptides were further diluted in PBS to the required concentrations.

### High throughput magnetic tweezers apparatus

Experiments were performed using a high throughput magnetic tweezers (MT) set-up previously detailed in Ref. (19, 18). In brief, it is a custom-built inverted microscope equipped with a 60x oil immersion objective. The position of the objective is controlled by a PIFOC piezo (P-726, Physik Instrumente Germany). A flow cell is mounted on the objective and illuminated using a collimated LED-light beam. A CMOS camera set to an acquisition frequency of 58 Hz and shutter time of 17 ms is used to image the field of view. A pair of 5 mm permanent magnet cubes (SuperMagnete, Germany) separated by a 1mm gap are positioned vertically above the flow channel. The vertical displacement and rotation of the magnets is controlled by linear motors, M-126-PD1 and C-150 (Physik Instrumente, Germany), respectively. The C++/CUDA bead tracking software used to monitor the beads has been extensively described in Cnossen et al. 2014 (21). It is open-source and available at http://www.github.com/jcnossen/qtrk. Additionally, a LabView measurement code integrated with the tracking software is available at http://www.github.com/jcnossen/BeadTracker. The temperature is controlled as described in Seifert et al., 2020 (27). In short, the objective is covered with a flexible resistive foil containing a 10 MΩ thermistor (HT10K, Thorlabs) and further insulated with Kapton tape (KAP22-075, Thorlabs). The temperature is regulated within ∼0.1°C by connecting the foil to a PID temperature controller (TC200 PID controller, Thorlabs).(21)

### Flow cell preparation

The flow cell assembly procedure has been described in Ref. (18). Briefly, a pre-cut double parafilm layer is sandwiched between two glass coverslips (#1, 24×60 mm, Menzel GmbH Germany). The bottom coverslip is coated with 5 μl of 0.3 mg/ml Nitrocellulose dissolved in amyl acetate and the top coverslip has a hole at each end, creating an inlet and outlet for the flow channel.

Upon mounting the flow cell on the MT objective, it is washed with phosphate-buffered saline (PBS) and incubated with 1 μm polystyrene reference beads (Sigma Aldrich). To functionalize the flow channel, 50 μl of 50 μg/ml of anti-digoxigenin is flushed into the flow channel and incubated for 30 min. Thereafter, the flow channel is rinsed with 1 ml of 10 mM Tris, 1 mM EDTA pH 8.0, 750 mM NaCl and 2 mM sodium azide buffer to wash out any excess anti-digoxigenin. Following a rinse with PBS, the flow channel is passivated with 90 μl of 10 mg/ml of bovine serum albumin (BSA, Sigma Aldrich) for 30 min to prevent non-specific binding to the flow cell.

To prepare the dsRNA tethers, 10 μl of Dynabeads MyOne streptavidin T1 magnetic beads (Thermofisher, Germany) were washed in 40 μl of PBS and subsequently mixed with coilable 3149 bp dsRNA and 1 μl of 100 mg/ml of BSA. The dsRNA tethers are flushed into the flow cell and incubated for approximately 3 min before washing out any excess or unbound dsRNA tethers. The tethers are then screened to select single tethers with the correct length for single molecule measurements (18).

### Magnetic tweezers experiments

To distinguish between torsionally constrained (coilable) and torsionally unconstrained (non-coilable)dsRNA tethers, rotation-extension experiments were performed by rotating the magnets from -30 turns to +30 turns at a speed of 1 turn/s at 4 pN. The end-to-end distance of a coilable tether decreases at 4 pN when rotated to +30 turns due to plectoneme formation, while remains unaffected when rotated to -30 turns due to the formation of melting bubbles (**Figure 1F**). In contrast, no change in extension is observed for non-coilable molecules when magnets are rotated from -30 to +30 turns (18). To observe the effect of the peptide on the twist of dsRNA, rotation-extension experiments were performed at a speed of 1 turn/s at a force of 4 pN and 0.5 pN before and after addition of the peptide on torsionally constrained tethers. Additionally, to determine the impact of the peptide on the contour and persistence length of dsRNA, three consecutive dynamic force-extensions were performed ranging from 8 pN to 0.1 pN at a speed of 0.3 mm/s before and after peptide incubation for torsionally unconstrained tethers. Furthermore, to investigate the environment-sensitive release mechanism of peptide 1’-1’ upon exposure to reducing conditions, the peptide was treated with either no TCEP or 1 mM of TCEP and incubated for ∼10 min before adding it to the flow cell.

All experiments were performed at 25°C and in PBS with 2 mM sodium azide. Upon completion of each experiment, the dsRNA tethers were cleared, and the flow cell was incubated in high salt buffer (TE 1X with 750 mM NaCl) for 10 min. Next, the flow cell was rinsed with PBS and new tethers were flushed in for the next experiment.

### Fitting force-extension data and measuring the change in dsRNA twist

The values for the contour and persistence length of dsRNA before and after the addition of the peptide were extracted by fitting a non-extensible worm-like chain (WLC) model (**Equation 1**) along the force-extension curves obtained for each tether in triplicates (22). The WLC model gives the force required to extend the RNA by an amount *L*_*ext*_, given its persistence length (*L*_*p*_) and contour length (*L*_*c*_),

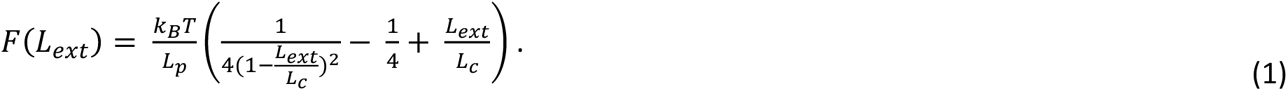

Here, *k*_*B*_ is the Boltzmann constant and *T* is the absolute temperature of the buffer. Given that at higher peptide concentrations the tethers collapsed at lower forces (**Figure S1A**), we only fitted the WLC along the first force-extension following peptide addition. However, due to large fluctuations in the position of the beads caused by Brownian motion, we integrated the force-extension curve to collapse the Brownian motion onto the curve (**Equation 2**)(20). This approach improves the estimation accuracy of the bead position during the force-extension curve.

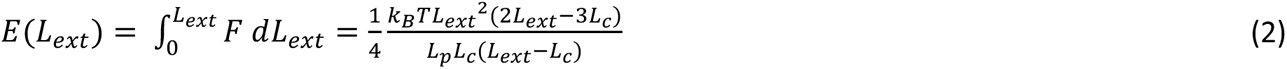

Individual force-extension curves were fitted with **Equation** 2, and the mean contour and persistence length across RNA molecules are reported. The standard deviations reported were calculated from the same data samples to estimate the error.

To observe the effect of the peptide on dsRNA twist, the change in extension recorded in the rotation extension experiments was plotted for each torsionally constrained dsRNA before and after the addition of the peptide. Next, to calculate the change in twist induced by the peptide, the maximum extension positions were manually selected for dsRNA before and after peptide addition. The calculated difference in peak positions provide a measure of the twist change induced by the peptide upon binding dsRNA in number of turns. To determine the slope of the plectonemic regions, we fitted a line along the rotation-extension traces between an extension of 0.2-0.4 μm.

### Fitting Hill model

To extract the dissociation constant (*K*_*d*_) and the Hill coefficient (*n*) for each peptide, we fitted the Hill model across the changes in twist with increasing peptide concentrations (*L*).

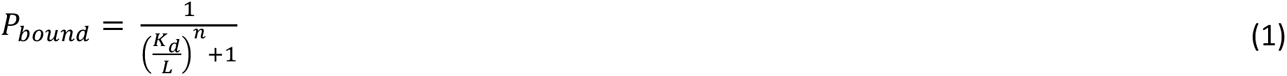

### Assuming the observed change in twist with respect to dsRNA (ΔΘ) is proportional to the fraction of RNA bound by the peptide (*P*_*bound*_), and that the RNA is fully covered by peptide at saturating conditions, we obtain

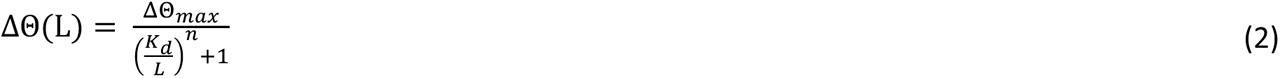

Fitting **Equation 2** to **Figure 2EG and 3C** allowed us to extract values for *n*, *n* and ΔΘ_*max*_. Error estimates represent 95% confidence intervals (± 2 SE) from 1000 bootstrap samples.

### Structural analysis of the TAV2b-dsRNA complex

To visualize alterations in the dsRNA helical structure upon interaction with TAV2b, we utilized USCF ChimeraX (28, 29). Using the ‘Matchmaker’ tool, we superimposed the TAV2b-dsRNA crystal structure (pdb: 2zi0) onto an A-form dsRNA of an identical sequence, generated using the ‘Build Structure’ tool. Next, we measured the distance across two nucleotides present in the major groove of the dsRNA, particularly using the phosphate atoms in the dsRNA backbone. These nucleotides were selected to measure the closest perpendicular distance between the backbone to obtain the shortest distance possible across the major groove. These measurements were recorded for both, free dsRNA and the dsRNA bound to TAV2b protein, to demonstrate dsRNA helical unwinding in the presence of TAV2b. Additionally, two base pairs marking a single helical turn were used to determine the number of base pairs present in each turn.

## Data availability

Link to be added.

## Supplementary Data

Supplementary Data are available online.

## Acknowledgements

DD and MK thank Jan Lipfert for useful discussion. DD and ZMR thank Samuel Schwab for assisting us with the structural analysis of the TAV2b-dsRNA protein complex.

## Funding

DD was supported by “BaSyC – Building a Synthetic Cell” Gravitation grant (024.003.019) of the Netherlands Ministry of Education, Culture and Science (OCW) and the Netherlands Organization for Scientific Research (NWO), and the NWO-M Open Competition Domain Science grant (OCENW.M.21.184).

## Conflict of interest statement

TNG is listed as an inventor in a patent for RNA-binding peptides.

## Authors Contribution

DD and TNG designed the research. ZMR, SQ and DD designed the experiments. ZMR performed the experiments and analyzed the data. MK and PAA helped with the Hill model and WLC fitting routines. QS made the dsRNA construct. TNG, MA, JP provided the peptide. ZMR and DD wrote the article. All the authors edited the article.

## Notes

### Competing Interest Statement

Tom N. Grossmann is listed as an inventor in a patent for RNA-binding peptides.

