## Supplementary Information for "Single-molecule magnetic tweezers reveals that TAV2b-derived peptides underwind and stabilize double-stranded RNA"

The Supplementary Information contains 3 Supplementary Figures and 3 Supplementary tables.

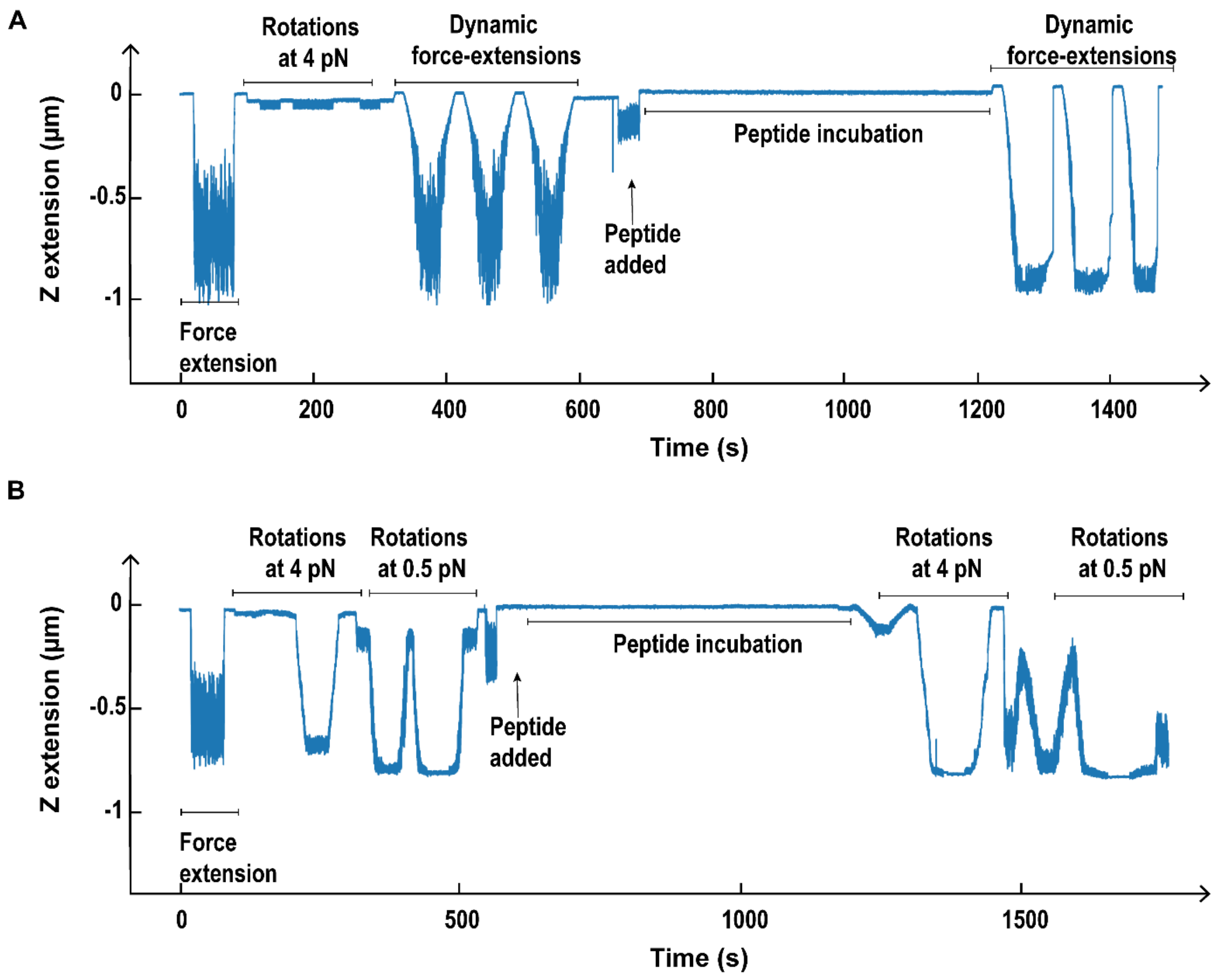


**Figure S1: Annotated raw experimental traces highlighting the assay performed with magnetic tweezers for peptide wt33. (A)** Time trace reporting on the tether extension when performing successive dynamic force extension assays. Trace is from experiments performed with 10 µM of peptide wt33 and is used for representation purposes. **(B)** Time trace reporting on the tether extension when performing successive rotation extension assays at 0.5 pN and 4 pN. The trace is from experiments performed with 8 µM of peptide wt33 and is used for representation purposes. (**A, B**) Peptide is added and incubated at 8 pN.

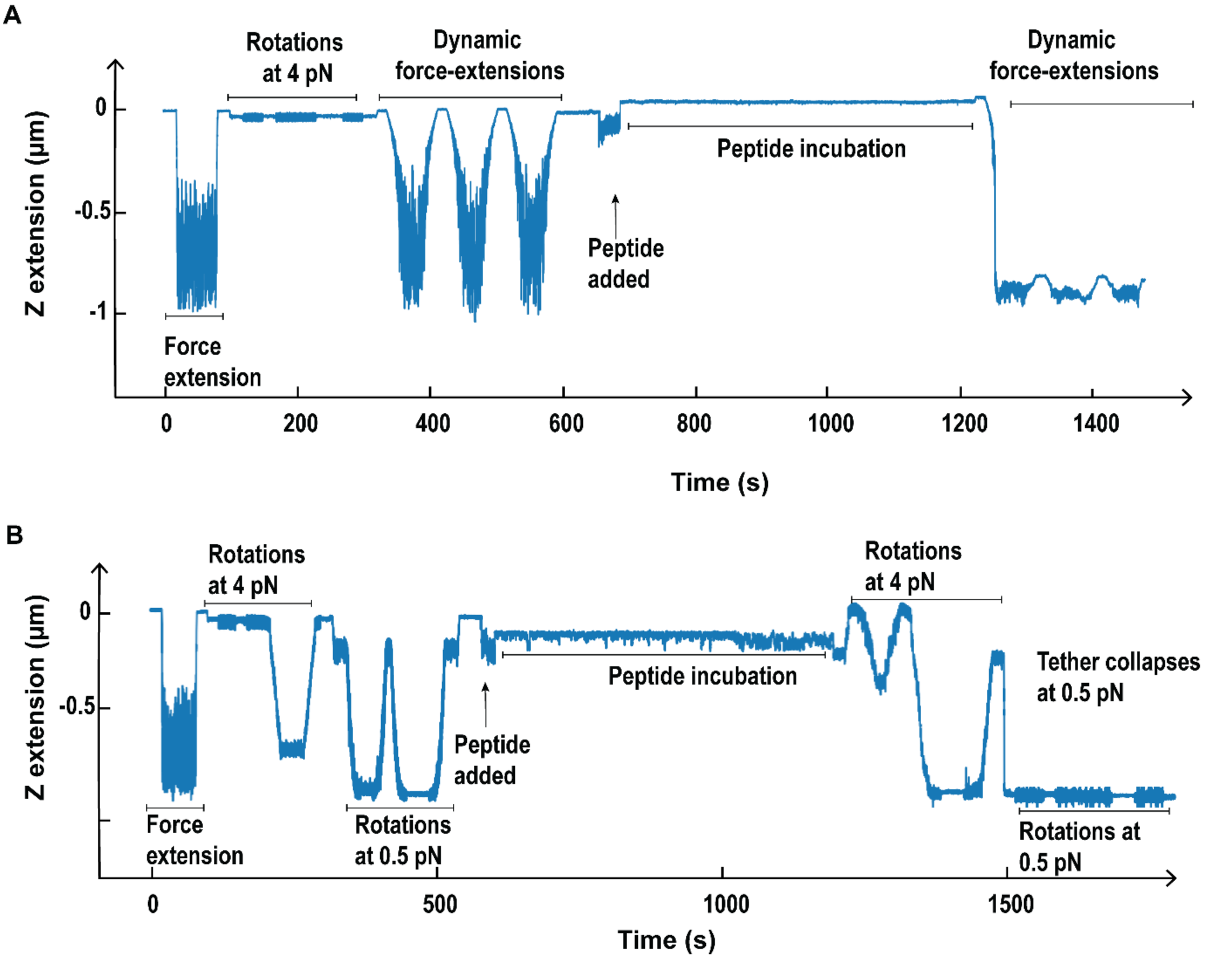


**Figure S2: Annotated raw experimental traces highlighting the assay performed with magnetic tweezers for peptide 1'-1'. (A)** Time trace reporting on the tether extension when performing successive dynamic force extension assays. Trace is from experiments performed with 7 µM of peptide 1'-1' and is used for representation purposes. **(B)** Time trace reporting on the tether extension when performing successive rotation extension assays at 0.5 pN and 4 pN. The trace is from experiments performed with 6 µM of peptide 1'-1' and is used for representation purposes. (**A, B**) Peptide is added and incubated at 8 pN.

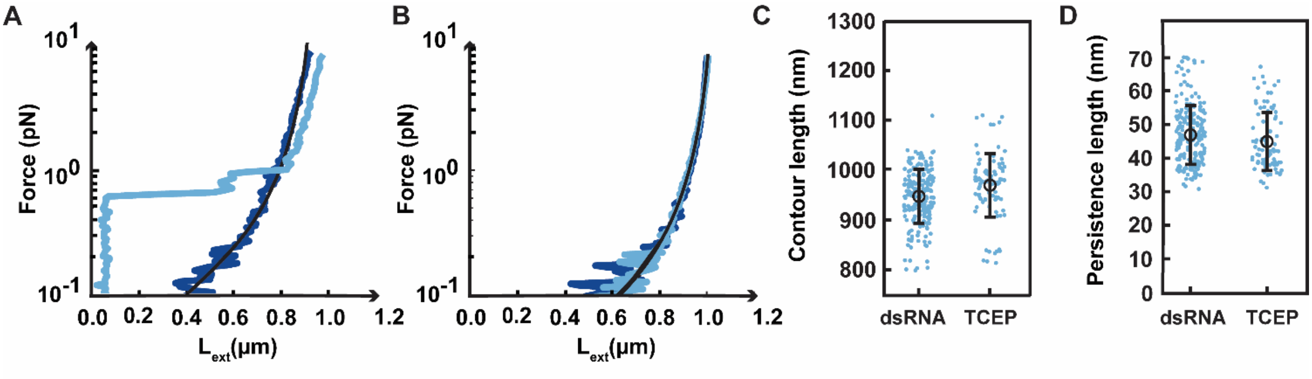


**Figure S3: Peptide 1′-1′ cannot bind to dsRNA upon exposure to TCEP. (A)** Force-extension experiments for a torsionally unconstrained dsRNA tether before (dark blue) and after (light blue) the addition of 7 μM 1′-1′ peptide. **(B)** Force-extension experiments for a torsionally unconstrained dsRNA tether before (dark blue) and after (light blue) the addition of 7 μM 1′-1′ peptide treated with TCEP. The black solid line is a WLC model fit to the data. **(C)** Plot displaying the distribution of contour length for torsionally unconstrained dsRNA with 7 μM of peptide 1′-1′ treated with TCEP. **(D)** Plot displaying the distribution of persistence lengths for torsionally unconstrained dsRNA with 7 μM of peptide 1′-1′ treated with TCEP.

**Table S1: Mean contour and persistence length for dsRNA and in the presence of 10 µM wt33.**

| | Contour Length ± std | | | N | Persistence Length ± std | | | N |
|---|---|---|---|---|---|---|---|---|
| **dsRNA** | 968 | ± | 75 | 564 | 46 | ± | 9 | 564 |
| **wt33 (10 µM)** | 1092 | ± | 94 | 70 | 10 | ± | 2 | 70 |

**Table S2: Mean induced twist (turns) on dsRNA upon binding of wt33 with increasing peptide concentrations measured at 0.5pN and 4 pN.**

| Concentration (µM) | Induced twist (turns) ± std At 0.5 pN | | | N | Induced twist (turns) ± std At 4 pN | | | N |
|---|---|---|---|---|---|---|---|---|
| **0.5** | -0.09 | ± | 0.4 | 46 | 0.02 | ± | 0.08 | 53 |
| **1** | -0.01 | ± | 0.4 | 47 | -0.1 | ± | 0.5 | 65 |
| **2** | -1.6 | ± | 0.8 | 55 | -1.2 | ± | 1.1 | 65 |
| **3** | -7.1 | ± | 1.1 | 55 | -4.5 | ± | 1.1 | 56 |
| **4** | -9.9 | ± | 1.4 | 63 | -5.7 | ± | 0.9 | 62 |
| **5** | -12.5 | ± | 1.3 | 66 | -7.2 | ± | 1.4 | 73 |
| **6** | -13.8 | ± | 2.0 | 31 | -8.0 | ± | 1.2 | 36 |
| **7** | -15.0 | ± | 1.5 | 30 | -8.6 | ± | 1.0 | 33 |
| **8** | -14.6 | ± | 1.3 | 54 | -8.8 | ± | 1.2 | 63 |
| **10** | - | - | | - | -9.9 | ± | 1.1 | 65 |

**Table S3: Mean induced twist (turns) on dsRNA upon binding of peptide 1'-1' with increasing peptide concentrations measured at 4 pN.**

| Concentration (µM) | Induced twists (turns) ± std | | | N |
|---|---|---|---|---|
| **0.5** | -0.4 | ± | 0.9 | 45 |
| **1** | -3.1 | ± | 1.0 | 17 |
| **2** | -9.6 | ± | 2.1 | 50 |
| **3** | 11.5 | ± | 1.8 | 58 |
| **4** | -13.4 | ± | 1.5 | 28 |
| **5** | -13.1 | ± | 1.8 | 64 |
| **6** | -13.3 | ± | 1.8 | 45 |
| **7** | -13.0 | ± | 1.9 | 29 |
